# TRIO-AI: Hybrid temporal graph, ODE, and VAE modeling for high-resolution cellular trajectory inference in liver injury

**DOI:** 10.64898/2025.12.17.694956

**Authors:** Hui Li, Jinlian Wang, Yankai Wen, Hongfang Liu, Cynthia Ju

## Abstract

Resolving dynamic cellular transitions at single-cell resolution is essential for understanding complex biological processes in development, disease, and regeneration. However, existing trajectory inference methods struggle to capture heterogeneous temporal dynamics, particularly for rare or transitional cell populations critical to injury and repair responses. Here, we present TRIO-AI, a hybrid computational framework that synergistically integrates three complementary approaches: Temporal Graph Neural Networks (Temporal GNN) for branching detection, Neural Ordinary Differential Equations (Neural ODEs) for continuous state flow modeling, and Time-Variational Autoencoders (Time-VAE) for density-based transitional state identification.

We applied TRIO-AI to single-cell RNA sequencing data from mouse liver ischemia-reperfusion (IR) injury across multiple timepoints (days 0, 1, 3, and 5 post-injury), comprehensively characterizing major hepatic macrophage populations. TRIO-AI identified a distinct transitional macrophage population (MPs_3) that peaks at 48 hours post-injury and exhibits coordinated signatures for lipid handling (LRP1, LRP6, ABCA1, LDLR, SCARB1), efferocytosis (MARCO, MERTK, MSR1), and extracellular matrix engagement (ITGA9, ITGB1, SDC2). This population serves as a critical integrator of multicellular pro-reparative signals through APOE-LRP1 and FN1-ITGA9 signaling axes.

Spatial transcriptomics validation confirmed MPs_3 localization to peri-necrotic borders with dramatic enrichment of APOE-LRP1 co-localization at 48 hours (19 pairs) compared to 24 hours (3 pairs). Comparative analysis against five state-of-the-art methods demonstrated TRIO-AI’s superior performance in detecting transitional states, accurately reconstructing branching trajectories, and identifying off-path populations missed by conventional approaches. This framework provides a powerful platform for dissecting cellular dynamics in complex biological systems and reveals new insights into macrophage-mediated tissue repair mechanisms.

## Introduction

Ischemia-reperfusion (IR) injury remains a major cause of acute and chronic liver damage, with macrophages serving as central mediators of the hepatic immune response during both inflammatory and tissue repair phases [1,2]. Traditional classification schemes that distinguish only between resident Kupffer cells and monocyte-derived macrophages fail to capture the full complexity and functional heterogeneity of the macrophage response [3,4]. Single-cell RNA sequencing (scRNA-seq) combined with advanced trajectory inference now enables high-resolution dissection of macrophage dynamics at cellular and molecular levels, providing opportunities to identify novel subpopulations based on their transcriptional programs, lineage trajectories, and temporal activation states [5–7].

Understanding cellular trajectories in dynamic biological systems is central to deciphering complex processes such as development, regeneration, and disease progression. Such insights provide candidate markers for early diagnosis, prognosis, and monitoring of treatment response in liver IR injury [8]. While numerous computational tools have been developed to infer these trajectories, each exhibits unique strengths but inherent limitations that become apparent when modeling the full complexity of temporal single-cell data [9–11].

Widely used trajectory inference tools such as Monocle3 [12] and Slingshot [13] rely on static pseudotime assumptions and often fail to capture the true dynamic nature of cell transitions, particularly in modeling non-monotonic or reversible trajectories increasingly recognized in biological systems. RNA velocity-based methods (scVelo) [14] and probabilistic frameworks (CellRank) [15] introduced directionality using spliced and unspliced mRNA counts, representing a major advance. However, these approaches still operate on snapshot-based assumptions without integrating actual timepoint data or underlying cell-cell interaction graphs, resulting in limited temporal resolution and predictive power.

Methods that incorporate timepoints, such as Tempora, perform inference at the cluster level, thereby losing critical single-cell granularity necessary to detect transitional or rare cell states [16]. Dynamic optimal transport-based approaches (TrajectoryNet, TIGON) [17] leverage mathematical frameworks to infer trajectories but lack biological specificity, relying on generic geometric priors rather than domain knowledge such as cell-cell communication networks. Static clustering methods including Leiden and Louvain are agnostic to temporal structure and systematically miss rare, off-path cell populations that play key roles in disease or repair processes [18,19].

To address these limitations, we present TRIO-AI, a hybrid computational framework that synergistically integrates Temporal Graph Neural Networks for branching detection, Neural Ordinary Differential Equations for continuous state flow modeling, and Time-Variational Autoencoders for latent density-based transitional state refinement. This multi-level approach captures both discrete lineage structures and continuous state transitions while maintaining sensitivity to rare and transitional cell populations.

## Methods

## 1. Experimental Design and Data Collection

### 1.1 Animal Models and Ethics

All animal procedures were approved by the Institutional Animal Care and Use Committee (IACUC) of UTHealth Houston (protocol AWC-23-0087) and conformed to NIH guidelines. Male C57BL/6J mice (12 weeks old) were maintained under specific pathogen-free conditions with ad libitum access to food and water under a 12-hour light/dark cycle. Liver ischemia-reperfusion surgery was performed as previously described.

### 1.2 Experimental Groups

Two complementary experimental cohorts were used to capture diverse hepatic cell populations: Experiment 1: Mice were harvested at day 0 (baseline), day 1, day 3, and day 5 post-IR (n=2 per timepoint). Single-cell suspensions were processed to enrich non-parenchymal hematopoietic populations while missing endothelial cells and resident Kupffer cells.

Experiment 2: Mice were harvested at day 0, day 3, and day 5 (n=2 per timepoint). Processing was optimized to enrich endothelial cells and macrophage subsets (including Kupffer cells), creating a complementary dataset.

### 1.3 Tissue Processing and Library Preparation

Liver nonparenchymal cells (NPCs) were isolated as previously described. Cell suspensions with >90% viability were labeled with sample tags using the BD Rhapsody Single-Cell Multiplexing Kit according to manufacturer instructions. Pooled labeled cells were loaded onto BD Rhapsody Cartridges and processed with the BD Rhapsody HT Xpress System. Whole transcriptome amplification (WTA) and sample tag libraries were generated using the BD Rhapsody WTA workflow. Libraries were quantified (Qubit) and quality-checked (Bioanalyzer) before pooling at recommended ratios for sequencing.

### 1.4 Public Dataset Integration

To validate trajectory dynamics and anchor early injury responses, we incorporated public single-cell liver IR datasets with explicit timepoints (GSE223561) [20]. This SuperSeries includes paired single-nucleus RNA-seq (snRNA-seq) and Visium spatial transcriptomics from human and mouse liver regeneration studies. Raw and processed matrices were retrieved from GEO, harmonized to our feature space, and jointly batch-corrected with in-house datasets.

## 2. Computational Analysis Pipeline

### 2.1 Quality Control and Preprocessing

Raw single-cell count matrices underwent standard quality control filtering, removing cells with high mitochondrial transcript fractions (>20%), low gene counts (<200), or excessive total counts indicative of doublets. After QC, datasets from all timepoints were concatenated and normalized jointly using library size normalization followed by log-transformation. Highly variable genes were identified using the Seurat variance-stabilizing transformation method. Principal component analysis (PCA) provided initial dimensionality reduction, followed by batch correction using Harmony when integrating multiple experiments or public datasets.

### 2.2 Cell Type Annotation

Neighborhood graphs were constructed using the first 30 principal components, and cell clustering was performed using the Leiden algorithm at multiple resolutions. Cell type annotation employed a hybrid strategy combining manual biomarker-based identification and automated enrichment-based annotation. Cluster-specific marker genes were identified using the Wilcoxon rank-sum test. Canonical biomarkers for major liver cell types were used to assign initial identities. Macrophage populations were further refined into resident Kupffer cells and monocyte-derived macrophages based on established markers.

### 2.3 TRIO-AI Framework Architecture

The TRIO-AI framework integrates three complementary computational modules into a unified temporal-graph modeling pipeline. Each component addresses specific limitations of existing methods while synergistically enhancing overall trajectory inference capability. See Figure 1 Supplementary Table 1.

**Figure 1.**
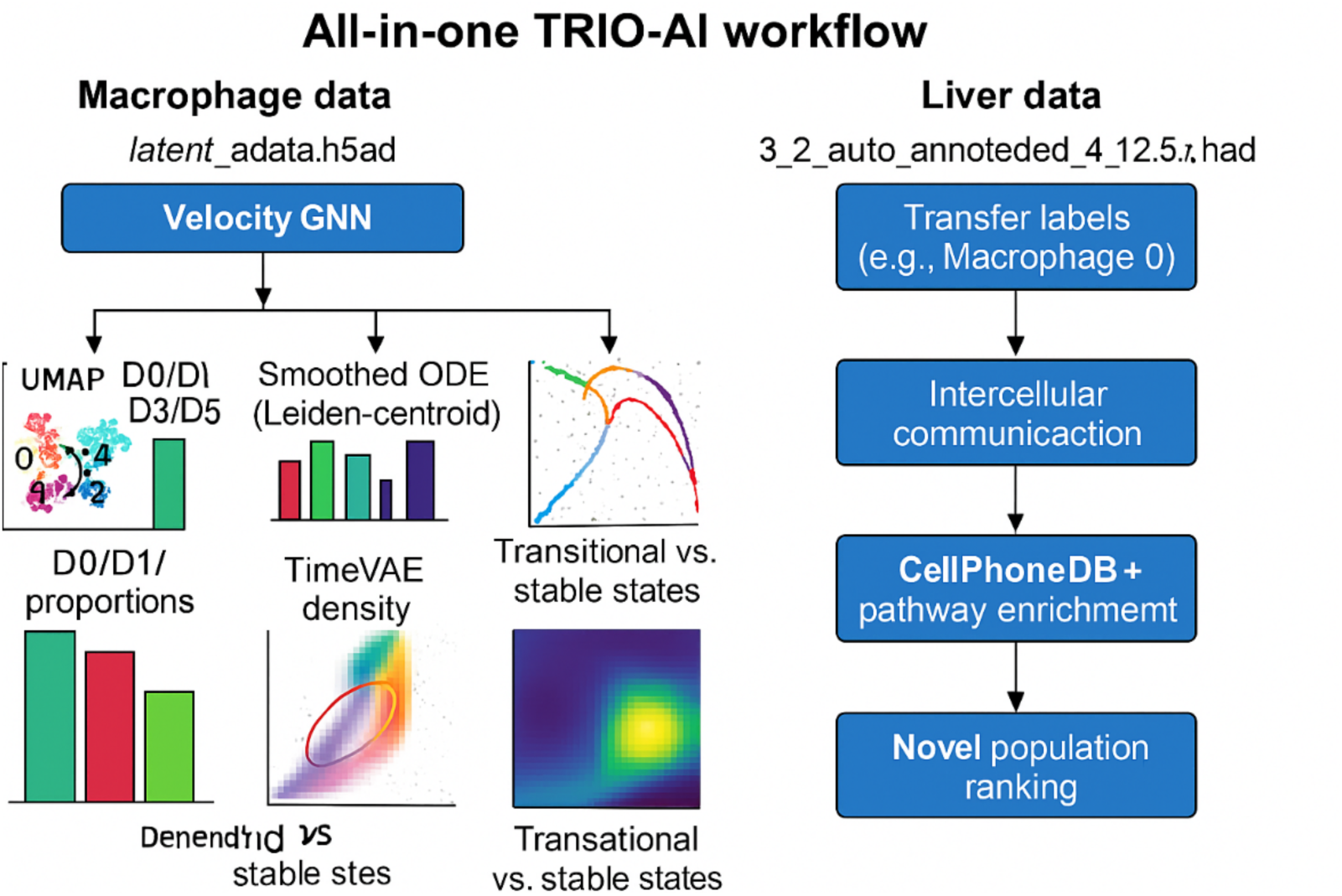
TRIO-AI Framework Architecture. Schematic diagram illustrating the integrated TRIO-AI framework showing data flow from raw expression through Temporal GNN embedding, Neural ODE trajectory modeling, and Time-VAE density analysis. Arrows indicate information flow between modules.

### 2.4 Temporal Graph Neural Network (GNN) Layer

The GNN layer constructs a dynamic cell-cell graph G = (V, E) where nodes represent individual cells and edges encode transcriptional similarity, temporal relationships, or RNA velocity-derived directionality. We systematically evaluated multiple edge definitions including k-nearest neighbors (kNN), mutual nearest neighbors (MNN), similarity thresholding, diffusion distances, PAGA cluster-level transitions, and directed RNA velocity graphs.

The Temporal GNN module processes single-cell data by constructing a feature matrix X from PCA-reduced gene expression (with optional diffusion pseudotime), z-score normalized, where nodes represent individual cells [21]. A k-nearest neighbor graph G = (V, E) is built with self-loops to preserve node identity. Temporal labels are derived by binning time_hours into discrete intervals, with optional continuous targets for regression.

The architecture consists of a Graph Attention Network (GAT) layer computing H₁ = ELU(GAT(X, E)) followed by dropout, with embeddings H saved to adata.obsm[“X_temporalGNN”]. A classification head produces logits for time bins, while an optional regression head predicts continuous time. Training optimizes a composite loss function combining weighted cross-entropy for time-bin classification, smooth L1 loss for temporal regression, and edge smoothness constraints. [22]

Comparative analysis revealed that GNN-derived embeddings using RNA velocity-informed edges combined with Leiden cluster connectivity produced superior alignment between latent time and experimental timepoints, with improved separation of macrophage subpopulations and more coherent branching trajectories corresponding to known biological phases of liver injury and repair.

### 2.5 Neural Ordinary Differential Equation (ODE) Layer

To reconstruct macrophage trajectories at different levels of granularity, we applied Neural ODE modeling to two complementary representations of the latent temporal manifold generated by the Temporal GNN).

First, cells were ordered by normalized latent time and partitioned into Nb=30 quantile-based temporal bins. For each bin, we computed its latent-space centroid. A minimum spanning tree (MST) was constructed between centroids to identify potential branch points. Neural ODEs were then trained to model continuous state transitions, where a neural network parameterized by θ governs the dynamics. Simulated trajectories were integrated using an ODE solver and projected onto UMAP space to visualize smooth progression between temporal bins.

Second, cells were aggregated by Leiden clusters, and we computed cluster centroids and mean latent time for each cluster. Clusters were ordered by latent time, and Neural ODEs were fit to these ordered centroids to generate trajectories aligned to discrete community structure. The UMAP projections from both approaches revealed coherent directional flow, with the Leiden-based view emphasizing major state transitions between transcriptionally distinct macrophage subsets, and the latent-time bin view capturing finer-grained intermediate states along the injury-repair continuum.

### 2.6 Time-Variational Autoencoder (Time-VAE) Density Layer

To detect over- and under-sampled regions in the temporal manifold, we implemented a timepoint-conditioned variational autoencoder using the scvi-tools framework. Expression counts were normalized to 10,000 counts per cell, log-transformed, and scaled. Each cell was assigned a numeric timepoint label to condition the model on experimental sampling time.(Sup Figure S2) The encoder maps input gene expression with timepoint label to a latent distribution. The decoder reconstructs expression from latent variables conditioned on time. Training minimizes the negative evidence lower bound (ELBO), balancing reconstruction accuracy and regularization.[23]

To quantify local cell density in latent space, we applied kernel density estimation (KDE) after scaling and projecting embeddings into two principal components. The resulting log-density scores were stored per cell and visualized on UMAP, enabling identification of high-density stable states and low-density transitional regions. Cells in the lowest quartile of latent density were designated as transitional states, potentially representing rare intermediates or unstable phenotypes.

### 2.7 Novel Population Identification Framework

To comprehensively identify novel and transitional macrophage populations during liver IR injury, we developed an integrated three-layer analytical framework combining the complementary strengths of Temporal GNN, Neural ODE, and Time-VAE modules. This systematic approach enables detection of rare populations missed by conventional methods through multiple orthogonal evidence streams: temporal dynamics, trajectory topology, and latent space density.

Layer 1 (Temporal Enrichment Analysis): We applied the Temporal GNN with RNA velocity-informed edges to generate refined cell embeddings integrating transcriptional states with temporal dynamics. Macrophages were re-clustered using Leiden community detection and visualized in UMAP space. For each cluster, we quantified relative abundance across experimental timepoints, identifying clusters with unusual temporal enrichment patterns—particularly those peaking transiently during intermediate timepoints rather than persisting across all phases.

Layer 2 (Trajectory Topology Analysis): We applied Neural ODE-based trajectory analysis at two resolutions: fine-grained latent-time bin level capturing subtle transcriptional transitions, and coarse-grained Leiden cluster level revealing major state transitions. Populations occupying critical topological positions were flagged as high-priority candidates, specifically: cells at trajectory branch points, cells along rare side paths diverging from main trajectories, and cells in transitional low-density regions.

Layer 3 (Density-Based Detection): The Time-VAE density layer computed local sampling probabilities in the learned latent space. Cells in the lowest density quartile (bottom 25% of kernel density estimates) were classified as “transitional” and examined for signatures of active state transitions, including heterogeneous co-expression of markers from multiple established clusters and enrichment for transcription factors regulating differentiation.

Populations identified by multiple independent methods received highest priority for experimental validation. This multi-method convergence substantially reduces false discovery rates while maintaining sensitivity for rare but biologically meaningful states.

### 2.8 Cell-Cell Communication and Spatial Validation

Cell-cell communication analysis was performed using CellChat on normalized expression data to infer ligand-receptor interactions across all cellular compartments. For macrophage subtypes including candidate novel populations, we quantified incoming and outgoing signaling strengths and identified significantly enriched pathways. Pathway enrichment analysis (Gene Ontology, KEGG, Hallmark) was performed on cluster-specific marker genes.

Spatial validation integrated snRNA-seq data with Visium spatial transcriptomics using an AI-assisted co-localization pipeline (cell2location + label transfer + proximity-constrained ligand-receptor mapping). APOE-centered interactions with macrophage receptors LRP1 were quantified near peri-necrotic borders across timepoints, with spatial proximity mapping confirming physical co-localization of ligand-receptor pairs.

### 2.9 Quantitative Validation of Novel Populations

To validate and prioritize candidate populations, we computed temporal dynamics metrics and trajectory-based novelty scores. Temporal metrics included: (1) Rarity (R = 1 - mean proportion), (2) Peak Prominence (PP), (3) Fold-Change versus baseline (FC = peak/D0 proportion), and (4) Resolution Drop (RD). Populations with high R, PP, FC, and RD were classified as dynamic transitional states.

Trajectory-based scoring integrated four features: (1) Low-Density score (LD), (2) Rarity score (RR), (3) Branchiness (BR), and (4) Velocity Alignment (VA). The composite novelty score (Novelty = 0.25×LD + 0.25×RR + 0.25×BR + 0.25×VA) ranked trajectories by likelihood of representing novel differentiation routes.

## Results

### 1. Dataset Characterization, Quality Control and Cell Type Annotation

Single-cell RNA-sequencing analysis of liver tissue from four timepoints (D0, D1, D3, D5) following ischemia-reperfusion injury yielded high-quality data after quality control filtering. Initial annotation identified major hepatic cell populations including hepatocytes, endothelial cells, cholangiocytes, hepatic stellate cells, and diverse immune populations (macrophages, neutrophils, T cells, B cells). Macrophage populations showed substantial compositional dynamics across the injury timecourse, with distinct patterns of expansion and contraction corresponding to inflammatory and resolution phases (**Figure 2**).

**Figure 2.**
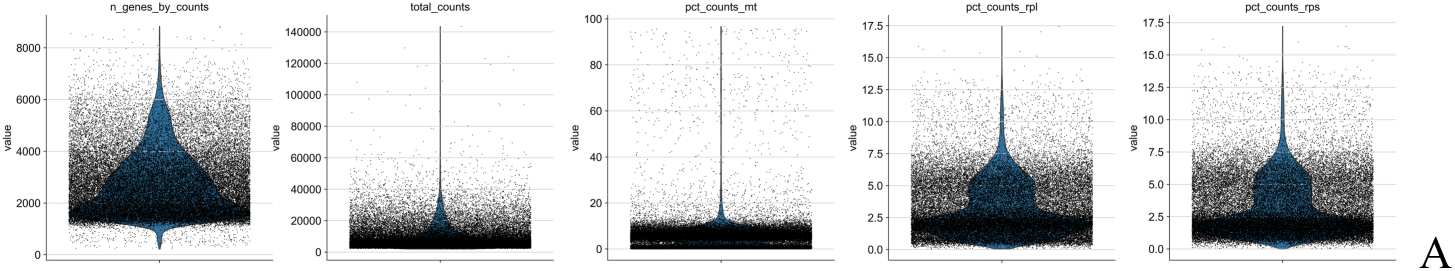
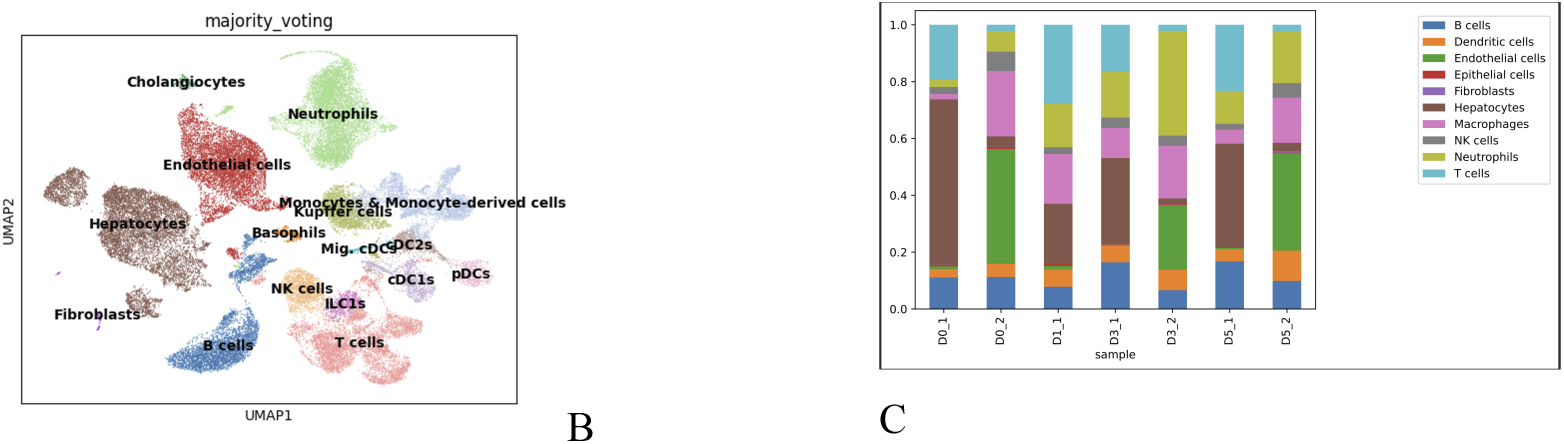
scRNA Data quality controls and cell type annotation. A. Data quality metrics, B. Cell types detected in the data set. C. Cell annotation.

### 2. TRIO-AI Framework Performance and Trajectory Reconstruction

Systematic comparison of embedding strategies revealed that UMAP projections based on Temporal GNN-derived features produced markedly improved temporal coherence compared to conventional PCA-based embeddings. Specifically, GNN embeddings using RNA velocity-informed edges demonstrated clear alignment between latent time gradients and experimental timepoints, with smooth progression from baseline through inflammatory to resolution phases. (**Figure 3).**

**Figure 3.**
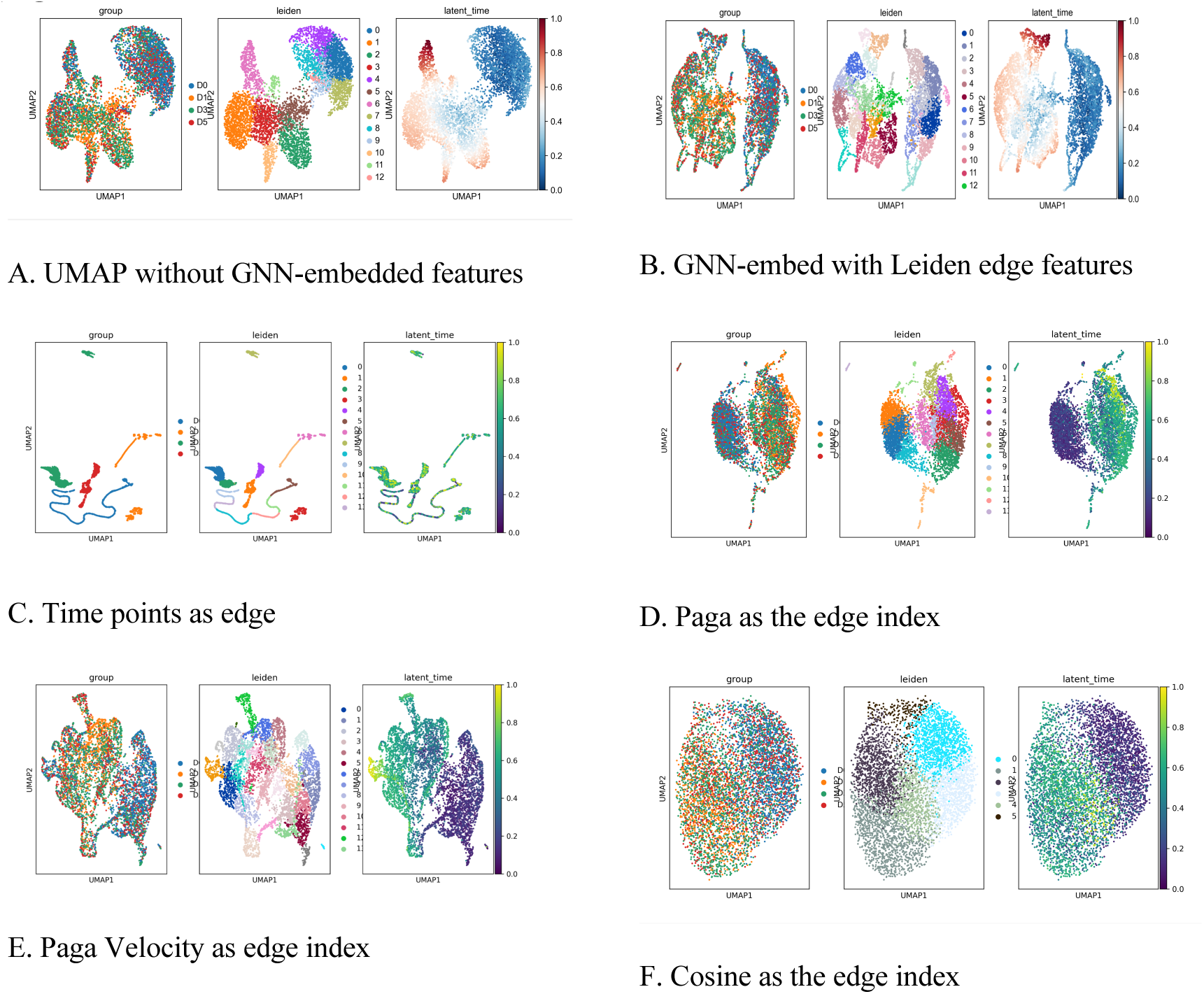
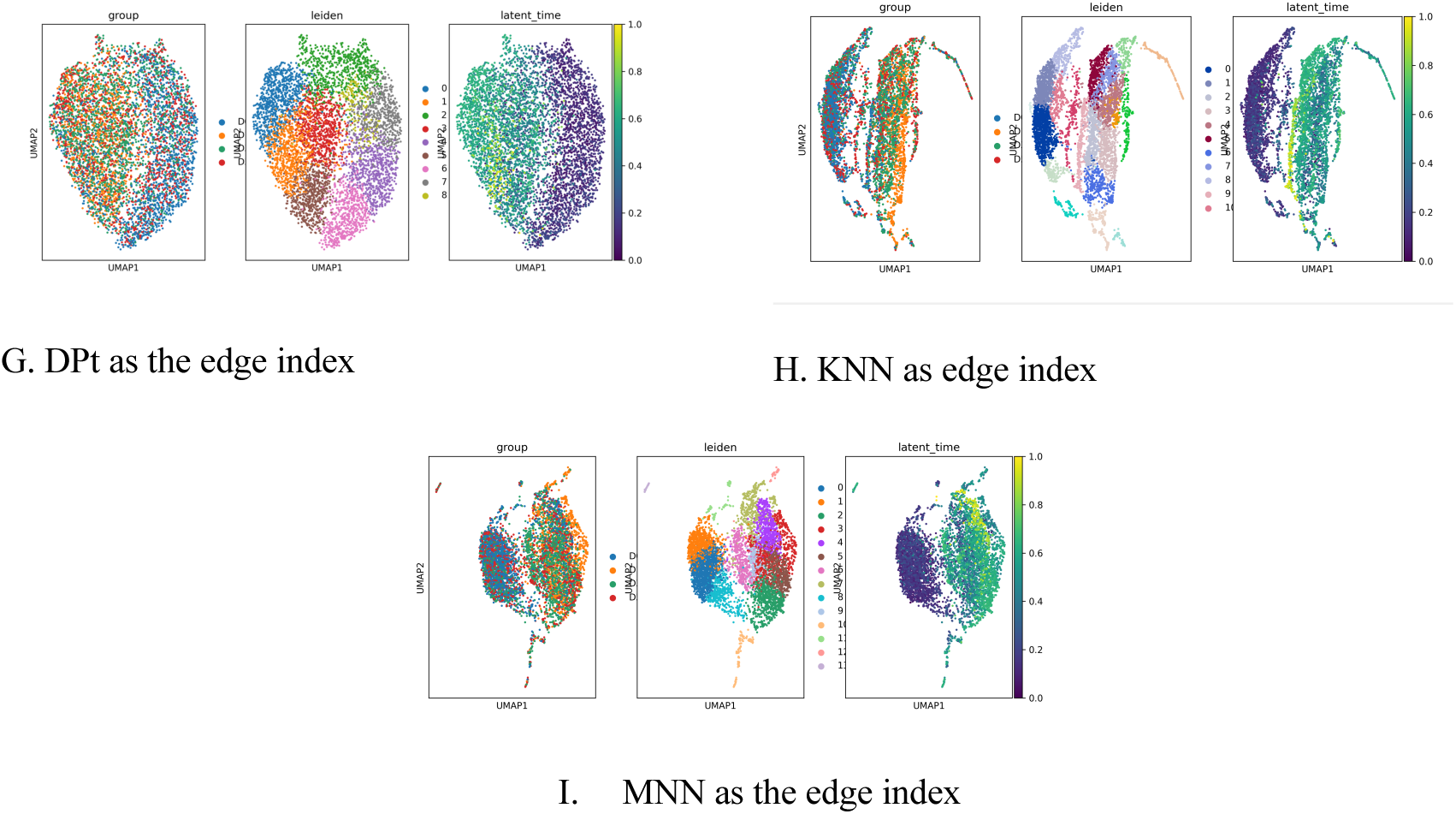
GNN Embedding Comparison. UMAP embeddings comparing (A) standard PCA-based representation, (B) GNN embedding with Leiden edge features, (C) timepoint-based edges, (D) PAGA-based edges, (E) velocity-based edges, (F) cosine similarity edges, (G) diffusion pseudotime edges, (H) kNN edges, (I) MNN edges. Color coding indicates experimental timepoints or latent time values.

Neural ODE trajectory reconstruction at both latent-time bin and Leiden cluster levels revealed coherent directional flow patterns. The bin-level view captured fine-grained intermediate states, while cluster-level trajectories emphasized major state transitions between transcriptionally distinct macrophage subsets. Integration of these complementary views provided comprehensive mapping of the macrophage state landscape.

Time-VAE density analysis identified distinct regions of high and low cell density in latent space, corresponding to stable attractor states and transitional regions, respectively. Low-density transitional zones localized between major macrophage clusters, suggesting the presence of intermediate or unstable phenotypes bridging distinct activation states. (**Figure 4**)

**Figure 4.**
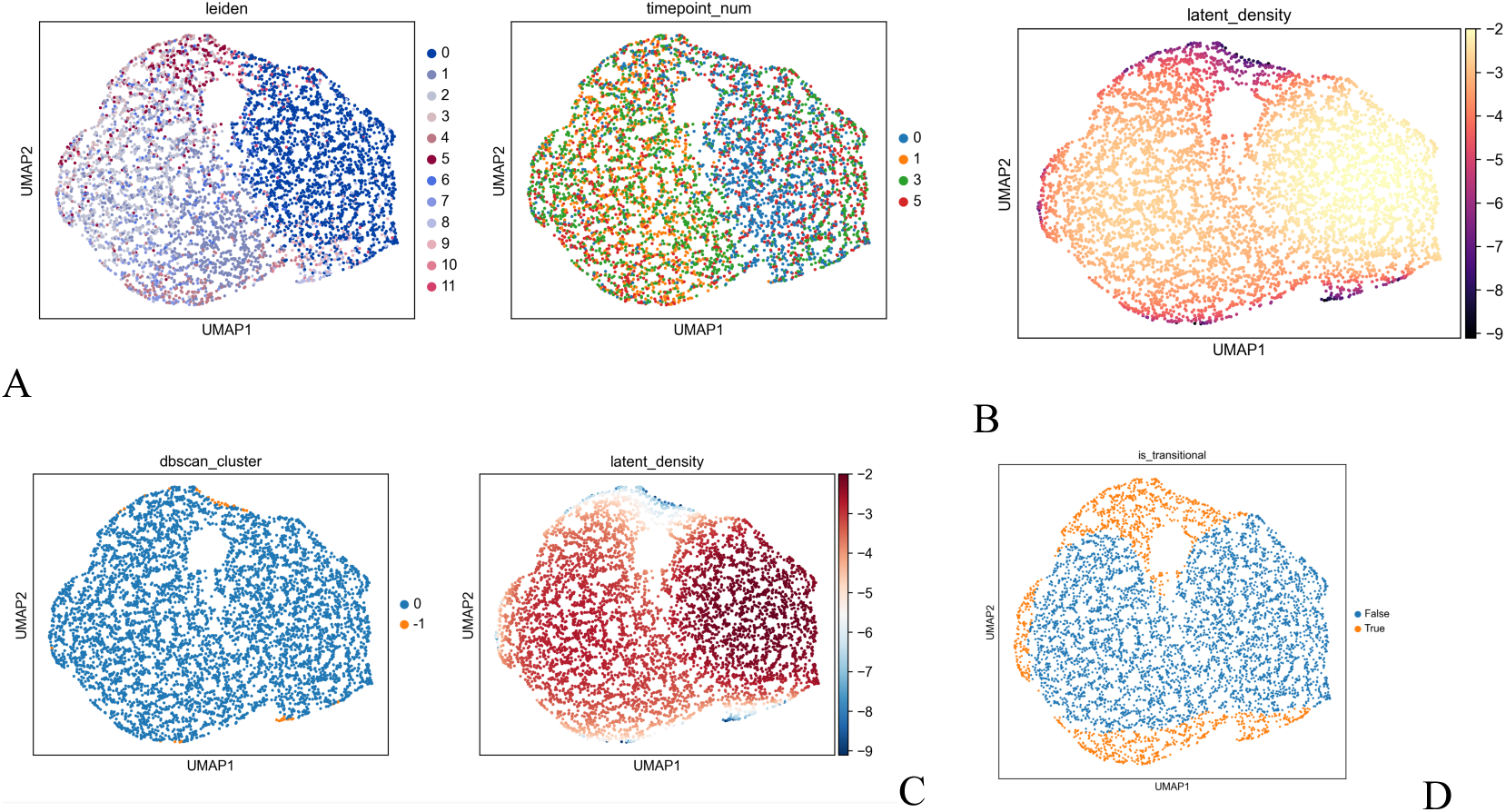
Time-Variational Autoencoder (Time-VAE) density analysis identifies transitional cell populations. UMAP embeddings colored by (A) experimental timepoints (D0, D1, D3, D5), (B) Leiden cluster assignments, (C) kernel density estimation (KDE) scores from Time-VAE, (D) individual cell sampling probabilities.

### 3. Identification of Novel Transitional Macrophage Population (MPs_3)

To identify novel macrophage populations during liver IR injury, we integrated trajectory dynamics, cluster-level temporal metrics, and graph-based topology analyses. Based on integrated temporal proportion analysis, trajectory positioning, and novelty scoring, clusters 3 and 7 emerged as strong candidates for novel macrophage subpopulations. Both clusters showed characteristic transitional dynamics: rare or absent at baseline (D0), sharp transient expansion at intermediate stages, and marked decline by D5. These patterns are consistent with short-lived, activation-dependent states bridging inflammatory and reparative phases.

Further integration and refinement in public dataset analysis (GSE223561) led to consolidation into a single high-confidence population designated MPs_3. This population exhibited peak abundance at 48 hours post-injury and occupied transitional regions in both Neural ODE trajectory space and Time-VAE density maps.(**Figure 5**)

**Figure 5.**
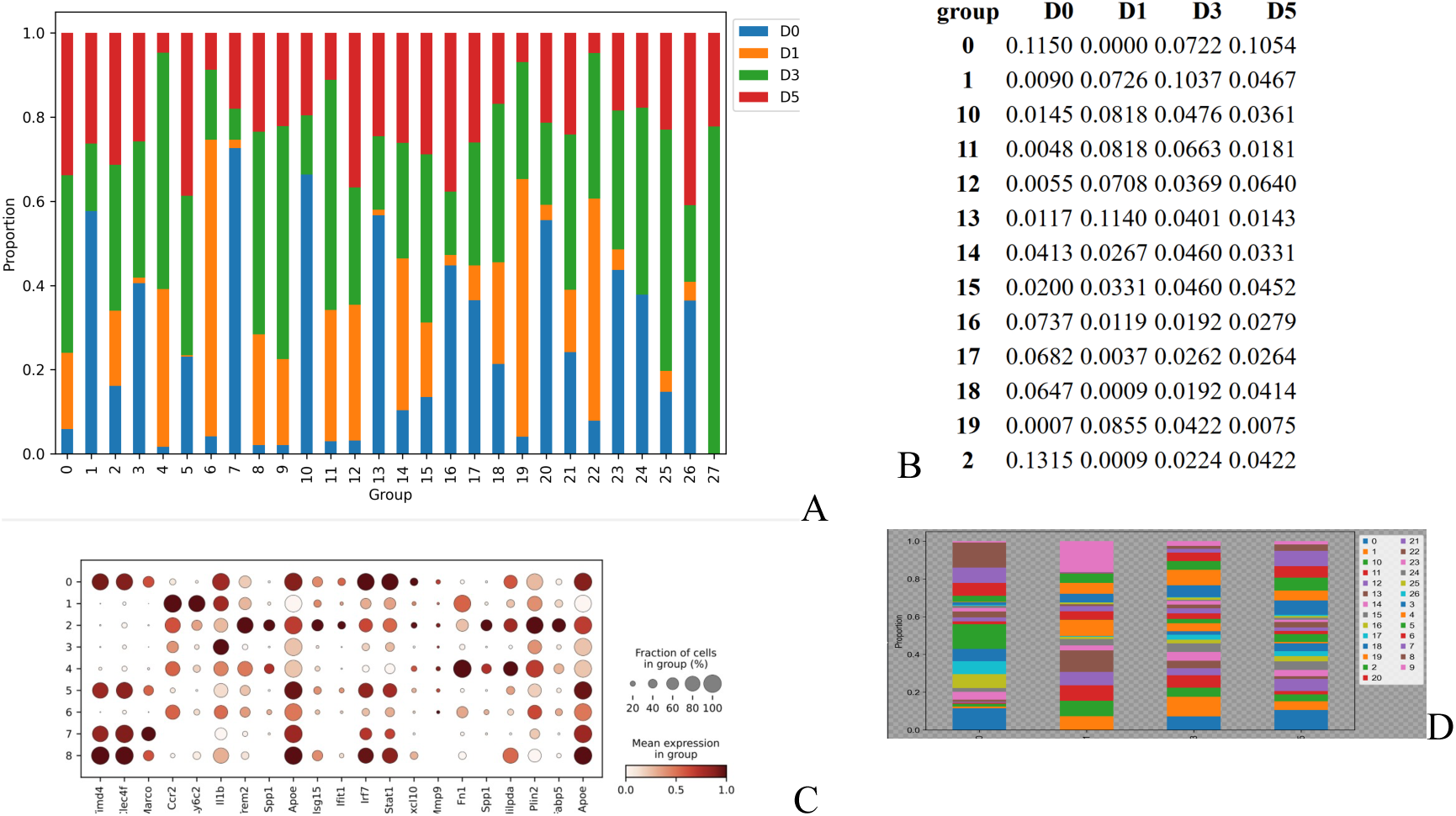
Cellular composition and transcriptional signatures across experimental conditions. **(A)** Stacked bar chart showing the relative proportion of cell clusters across individual samples or experimental conditions. Each color represents a distinct cell cluster, with vertical bars normalized to 100% to reveal compositional differences between samples. **(B)** Quantitative summary table listing cluster identities with corresponding statistical metrics, including cluster size, average gene counts, mitochondrial content percentages, and number of detected features per cluster. **(C)** Dot plot displaying marker gene expression across identified cell clusters. Dot size indicates the percentage of cells expressing each gene within a cluster, while color intensity represents average expression level (log-normalized). Row labels denote marker genes; column labels indicate cell clusters. (**D)** Compositional comparison across grouped conditions (e.g., control vs. treatment groups), demonstrating shifts in cell type proportions. Each colored segment represents a cell population identified through clustering analysis.

### 4. Transcriptional Characterization of MPs_3

Differential expression analysis identified a distinct transcriptional signature for MPs_3 characterized by coordinated programs for:

- Lipid handling and metabolism: LRP1, LRP6, ABCA1, LDLR, SCARB1, LPL, FABP5
- Core macrophage and reparative programs: TREM2, CX3CR1, CD68, SPP1, LYZ2
- Efferocytosis and scavenger functions: MARCO, MERTK, MSR1, LGALS3, GPNMB
- ECM engagement and integrin signaling: ITGA9, ITGB1, SDC2

Pathway enrichment analysis confirmed activation of PI3K-AKT, MAPK, and NF-κB signaling cascades, promoting macrophage survival, efferocytosis, and tissue repair functions.

Pseudotime trajectory analysis demonstrated progressive down-regulation of CCR2-associated inflammatory features and concurrent up-regulation of TREM2, SPP1, and cytoskeletal-ECM genes as cells transitioned through MPs_3, culminating in terminal reparative states. (**Figure 6**)

**Figure 6.**
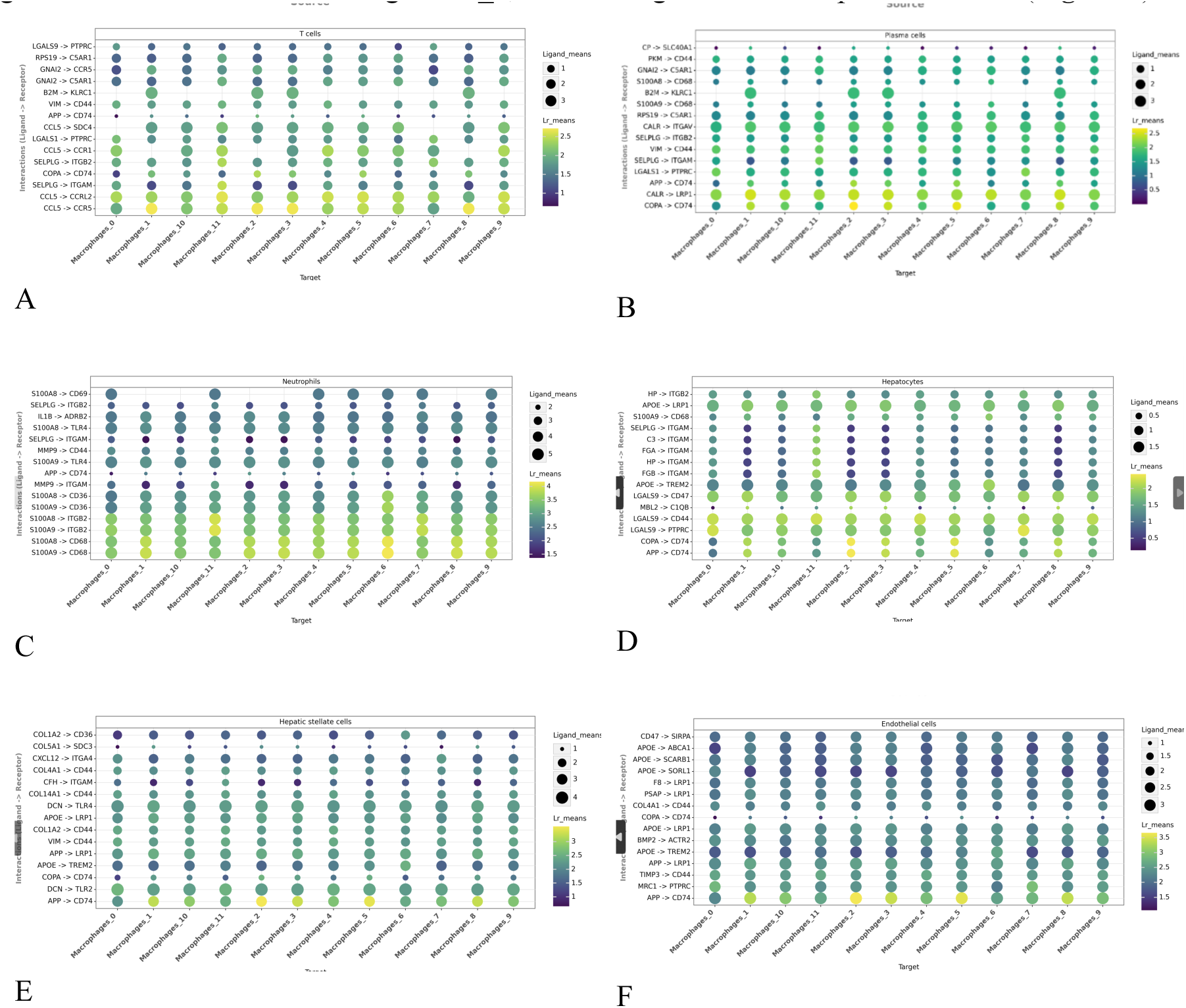
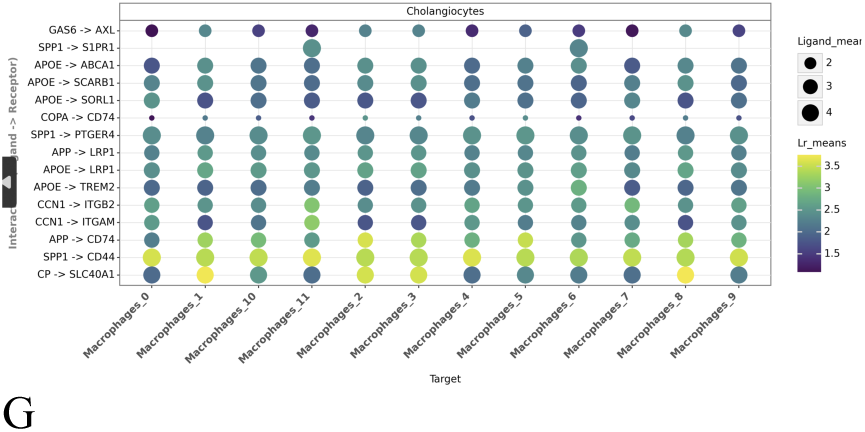
Multicellular signaling networks converge on MPs_3 macrophages during liver injury resolution. Comprehensive cell-cell communication analysis showing ligand-receptor interactions from diverse liver cell types to macrophage subpopulations. Each panel represents signaling from a specific cell type: (A) hepatocytes, (B) cholangiocytes (bile duct cells), (C) liver sinusoidal endothelial cells (LSECs), (D) hepatic stellate cells (HSCs), (E) B cells, (F) mesenchymal cells, and (G) other immune cells. Dot size represents mean ligand expression in source cells, while color intensity indicates ligand-receptor communication probability. MPs_3 (Macrophages_3) receives convergent signals across multiple cellular sources, particularly through apolipoprotein-mediated pathways (APOE, APOA1, APOB, APOC3 engaging LRP1, LRP6, ABCA1, TREM2), ECM ligands (FN1, PLG engaging ITGA9, ITGB1), and other factors (HP/HPX engaging LRP1, SDC2). This multicellular coordination establishes MPs_3 as a central integrator of tissue-wide repair signals. Analysis performed using CellChat (p < 0.05).

### 5. Cell-Cell Communication Networks Converging on MPs_3

Systematic ligand-receptor interaction analysis revealed MPs_3 as a major communication hub receiving diverse upstream signals. Key axes included apolipoprotein signaling (APOE→LRP1), ECM-integrin interactions (FN1→ITGA9), and heme-associated signaling (HP/HPX→LRP1). The APOE-centered network emerged as particularly prominent, with APOE engaging multiple receptors on MPs_3 (LRP1, LRP6, ABCA1), supporting integrated lipid handling. (**Figure 7**)

**Figure 7.**
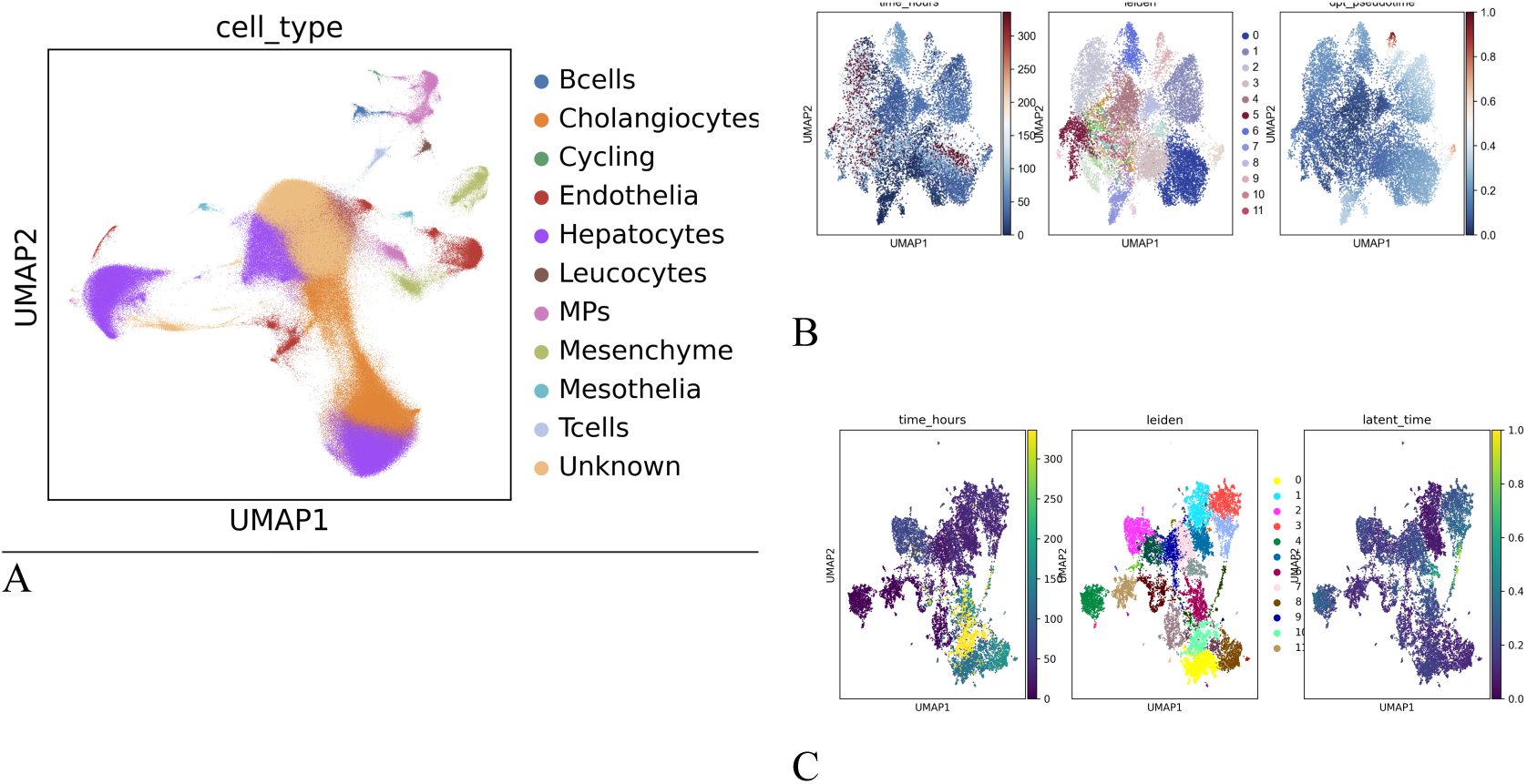
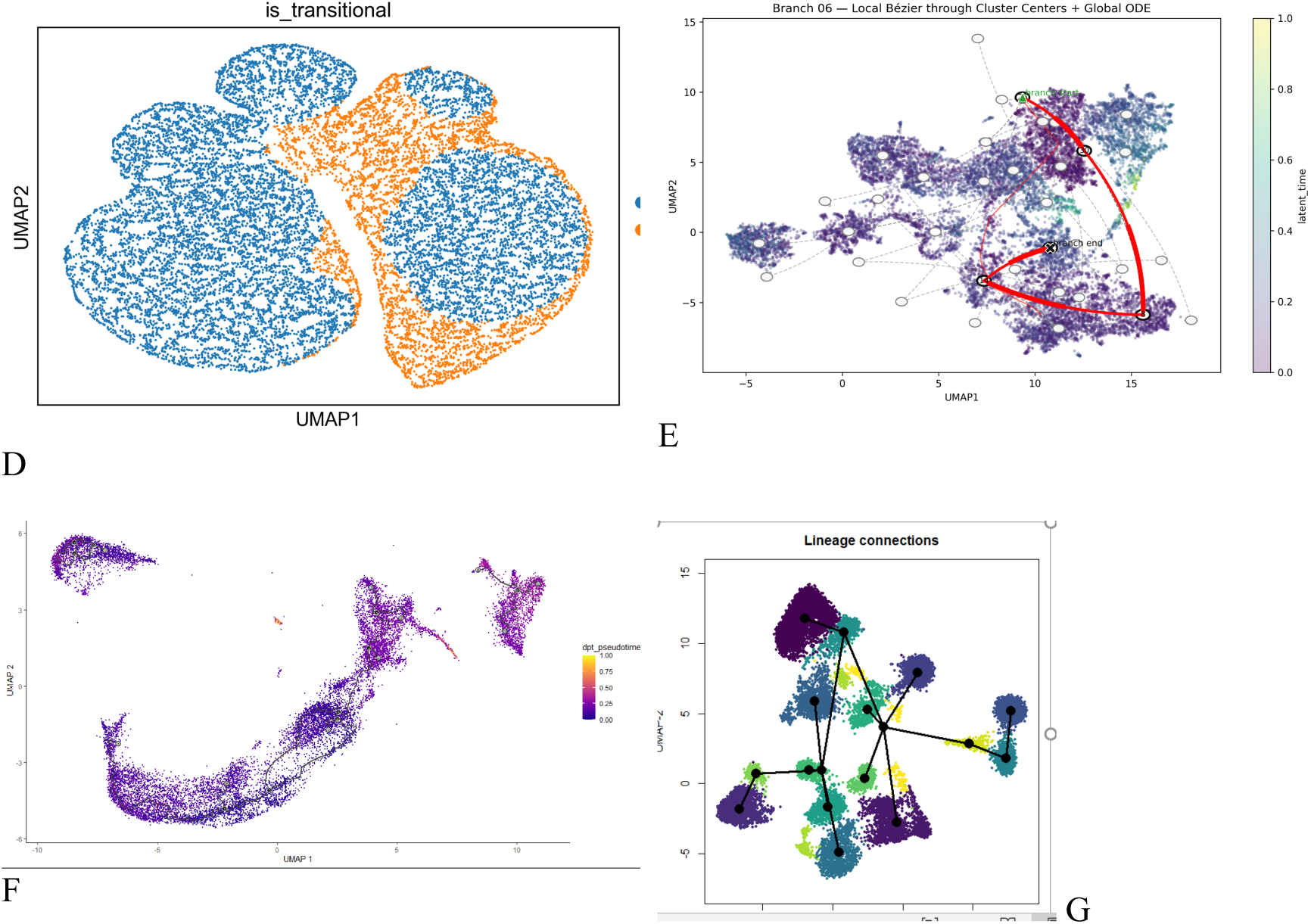
Temporal GNN embedding and hybrid trajectory inference reveal a reparative macrophage population. **(A)** UMAP visualization of single-cell transcriptomes colored by cell type, showing diverse immune and tissue-resident populations including macrophages, monocytes, T cells, B cells, and epithelial cells. **(B-C)** Expression patterns of key marker genes across the UMAP embedding, identifying cell type-specific signatures. **(D)** UMAP showing sample or condition distribution (blue/orange clusters) revealing batch effects or temporal states. **(E)** Trajectory analysis with directional flow indicating cellular state transitions and identification of a reparative macrophage population (highlighted region). **(F)** Pseudotemporal ordering of cells along differentiation trajectory, with color gradient representing progression through cellular states. **(G)** Lineage connectivity network derived from TemporalGNN analysis, illustrating relationships between cell populations and paths toward the reparative macrophage phenotype.

To investigate the diversity of macrophage populations, we performed comprehensive molecular profiling using single-cell transcriptomics. UMAP dimensionality reduction revealed multiple distinct macrophage subsets characterized by differential expression of key functional markers (Figure 8A). Inflammatory macrophages exhibited high expression of TNFAIP2 and SPP1, while tissue-resident macrophages were marked by MERTK and CD9. Matrix remodeling activity was indicated by expression of MMP1 and CHI3L1, and cellular activation states were defined by ITGAM and CD83 expression patterns.

**Figure 8.**
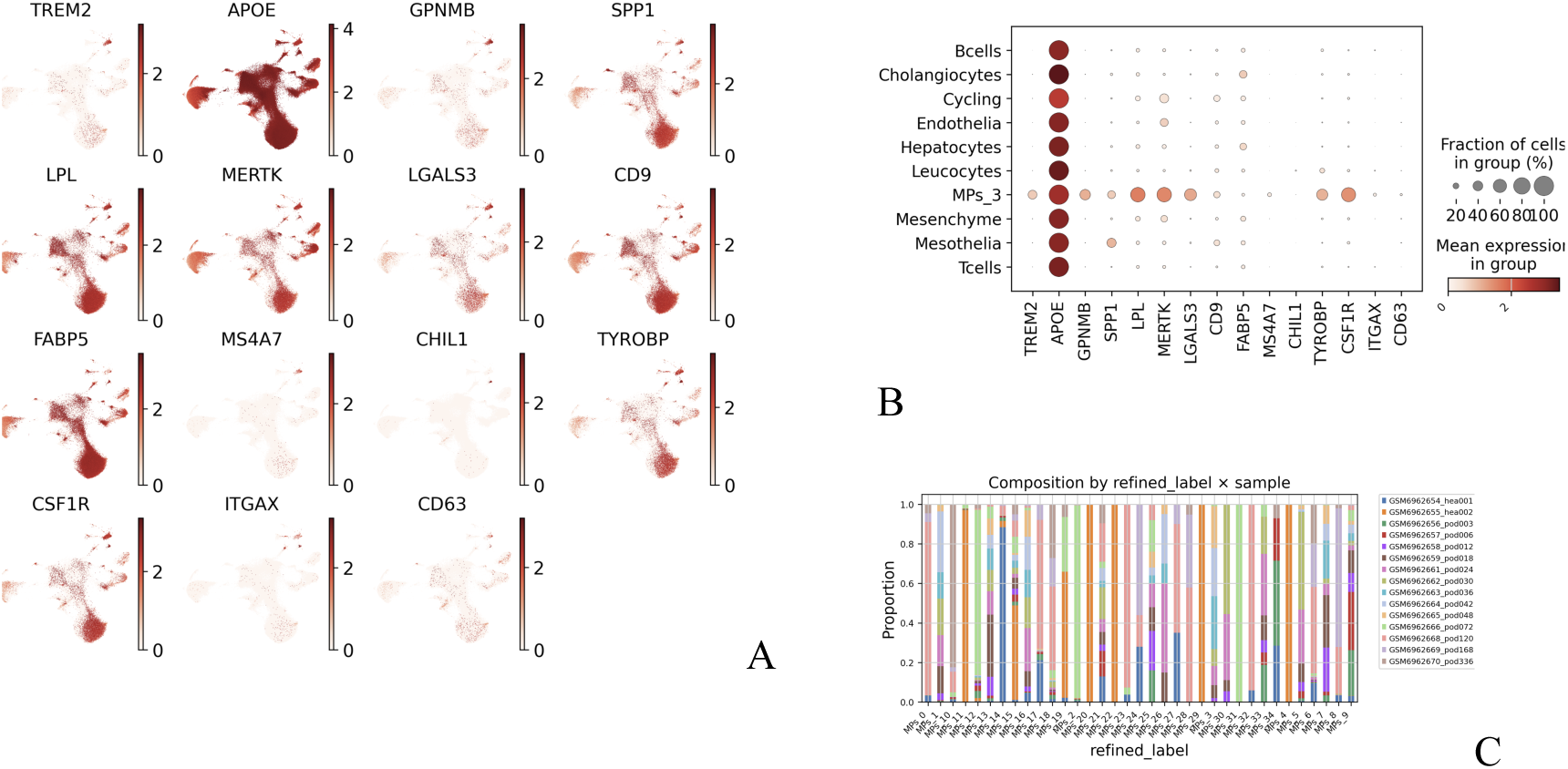
Molecular characterization and tissue distribution of macrophage subpopulations. **(A)** UMAP feature plots showing expression of key marker genes defining distinct macrophage states, including inflammatory markers (TNFAIP2, SPP1), tissue-resident signatures (MERTK, CD9), matrix remodeling genes (MMP1, CHI3L1), and activation markers (ITGAM, CD83). Expression intensity shown in red-brown gradient. **(B)** Dot plot depicting tissue-specific gene expression signatures across identified macrophage clusters. Dot size represents the percentage of cells expressing each gene; color intensity indicates average expression level. Tissue markers include brain, kidney, cartilage, placenta, lung, and spleen-specific genes. **(C)** Stacked bar chart showing the proportional contribution of each donor/sample to the defined macrophage states, revealing inter-individual variation in macrophage subset composition. Each color represents a distinct cellular state identified in panel A.

To assess tissue tropism and functional specialization, we examined tissue-specific gene expression signatures across the identified macrophage clusters (Figure 8B). Distinct macrophage subsets displayed enrichment for tissue-selective markers corresponding to brain, kidney, cartilage, placenta, lung, and spleen-specific genes, suggesting functionally specialized populations adapted to their respective tissue microenvironments. The percentage of cells expressing each marker gene and the average expression intensity varied substantially across clusters, highlighting the molecular diversity within the macrophage lineage.

Analysis of donor contribution to each macrophage state revealed considerable inter-individual variation in macrophage subset composition (Figure 8C). The proportional representation of distinct cellular states differed markedly between donors, with some individuals showing enrichment for specific macrophage phenotypes while others displayed more balanced distributions. This heterogeneity suggests that individual-specific factors—potentially including genetics, environmental exposures, or physiological state—influence macrophage population structure and functional polarization.

### 6. Spatial Validation and Temporal Dynamics

Integration with Visium spatial transcriptomics revealed dramatic temporal enrichment of APOE-centered intercellular interactions at 48 hours. APOE→LRP1 ligand-receptor pairs increased 6.3-fold (3 to 19 pairs). Analysis using CellChat identified CHI3L1 as a pivotal APOE receptor-complementing molecule mediating tissue damage resolution. These findings establish LRP1 as a parallel APOE receptor component in CHI3L1 co-expression networks, reinforcing its role in the transition to reparative macrophage states. (**Figure 9**)

**Figure 9.1.**
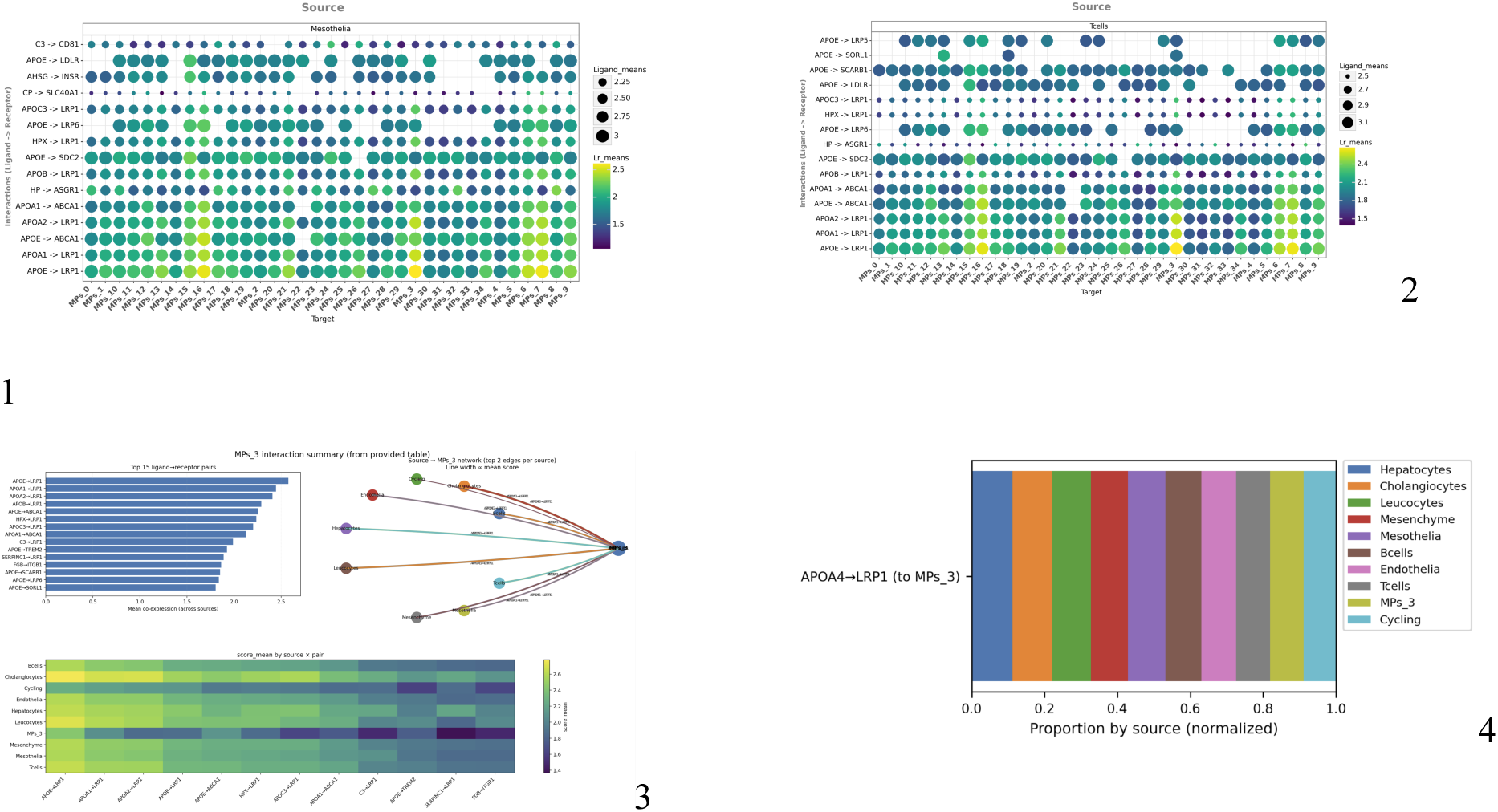
Ligand-Receptor Interactions from Non-Macrophage Sources to Macrophage Targets. Dot plot depicting prioritized L-R interactions where non-macrophage cell types serve as sources. Y-axis: Ligand genes (e.g., APOE, HPX); x-axis: Macrophage target subpopulations (e.g., MPs_3). Dot size represents interaction ranking (LIANA score); color scale indicates lr_means (log-normalized expression). MPs_3 receives the broadest and highest-confidence signals, with APOE-LRP1 emerging as a dominant pair (lr_means = 3.75). Figure 9.2. Ligand-Receptor Interactions from Non-Macrophage Sources to Macrophage Targets (Alternative View: Target-Centric). Dot plot visualizing the same L-R interactions, reoriented to emphasize macrophage targets on the x-axis and source cell types on the y-axis (e.g., Hepatocytes, Leucocytes). Dot size: Interaction ranking; color: lr_means (scale: 0–3.1). This view highlights source diversity, with hepatocytes and leucocytes as major contributors to MPs_3 signaling (e.g., APOE-SCARB1, lr_means = 2.4). Figure 9.3. Top L-R Interactions and Network Overview (A) Horizontal bar plot of the top 15 L-R pairs ranked by aggregate LIANA score, showing mean expression contributions from key ligands (e.g., APOE-APOC4). (B) Interaction network diagram illustrating directional signaling from source cell types (nodes colored by type: hepatocytes in blue, leucocytes in green) to macrophage targets (central hub). Edges represent high-confidence pairs, with thickness proportional to lr_means. This integrates ∼200 interactions, emphasizing APOE-centric hubs linking lipid-handling sources to reparative targets. Figure 9.4. Normalized Proportions of L-R Signaling by Source Cell Type Stacked bar plot quantifying the relative contribution of each source cell type to total incoming signals for the APOE-LRP1 interaction (a exemplar high-confidence pair). Bars represent normalized proportions (0–1) across macrophage targets (x-axis: MPs_1–MPs_3). Hepatocytes dominate (blue, ∼0.4–0.6), followed by leucocytes (green) and cholangiocytes (orange), underscoring multicellular coordination. Legend: Cholangiocytes (orange), Cycling (cyan), Endothelia (pink), Hepatocytes (blue), Leucocytes (green), Mesenchyme (gray), Mesothelia (brown), T cells (purple).

### 7. Integration of Cell-Cell Communication and Differential Expression Defines a Reparative Macrophage Signature

Building on ligand-receptor (L-R) interaction networks and differential gene expression (DEG) profiling, we established a high-confidence molecular signature for the transitional reparative macrophage subpopulation (MPs_3). By prioritizing LIANA-derived L-R pairs and intersecting them with upregulated DEGs, followed by KEGG pathway enrichment analysis, we identified a core functional program centered on lipid handling, efferocytosis, and extracellular matrix (ECM) remodeling (**Figure S1**).

A ranked marker-conviction table (top 20 shown; full dataset in *mps3_marker_conviction_table.csv*) highlights these convergent signals, combining three evidence streams: **(1)** APOE, LRP1, and SCARB1 (lipid import/efflux); **(2)** TREM2, CX3CR1, and CD68 (core macrophage and reparative functions); **(3)** MERTK, MSR1, MARCO, LGALS3, and GPNMβ (efferocytosis and scavenger activity); and **(4)** ITGA9, SDC2, and ITGB1 (ECM/integrin interactions). Notably, many of these markers are cell-surface-exposed and align with L-R interaction partners (e.g., LRP1–APOE; TR1B1, including the APOE→LRP1/LRP6/ABCA1 and PNI/PLG→ITGA9 pathways). This molecular convergence is reinforced by KEGG enrichment analysis, with key pathways including “ECM-receptor interactions,” “Fc-γ-mediated phagocytosis,” and “ECM-receptor interactions,” contrasted with downregulated terms such as “fatty acid and PPAR metabolism” in non-MPs_3 regions.

Pseudotime trajectory analysis further contextualizes this signature. MPs_3 progressively downregulates inflammatory markers (early trajectory branches 1–13) while coordinately upregulating lipid handling, efferocytosis, and ECM remodeling modules, culminating in repair-zone endpoints (trajectory branches 14–17). This temporal progression mirrors the observed activation-to-resolution arc, positioning MPs_3 as a dynamic integrator of reparative cues in tissue repair microenvironments.

We did KEGG enrichment analysis of differentially expressed genes in MPs_3 reveals a shift from inflammatory (e.g., chemokine signaling, upregulated) to reparative programs (e.g., ECM-receptor interaction, Fcγ-mediated phagocytosis), with suppression of metabolic catabolism (e.g., fatty acid degradation, PPAR; adjusted p<0.01). This aligns with MPs_3’s lipid-handling and efferocytosis signature, as visualized across subpopulations.(**Figure 10**)

**Figure 10.**
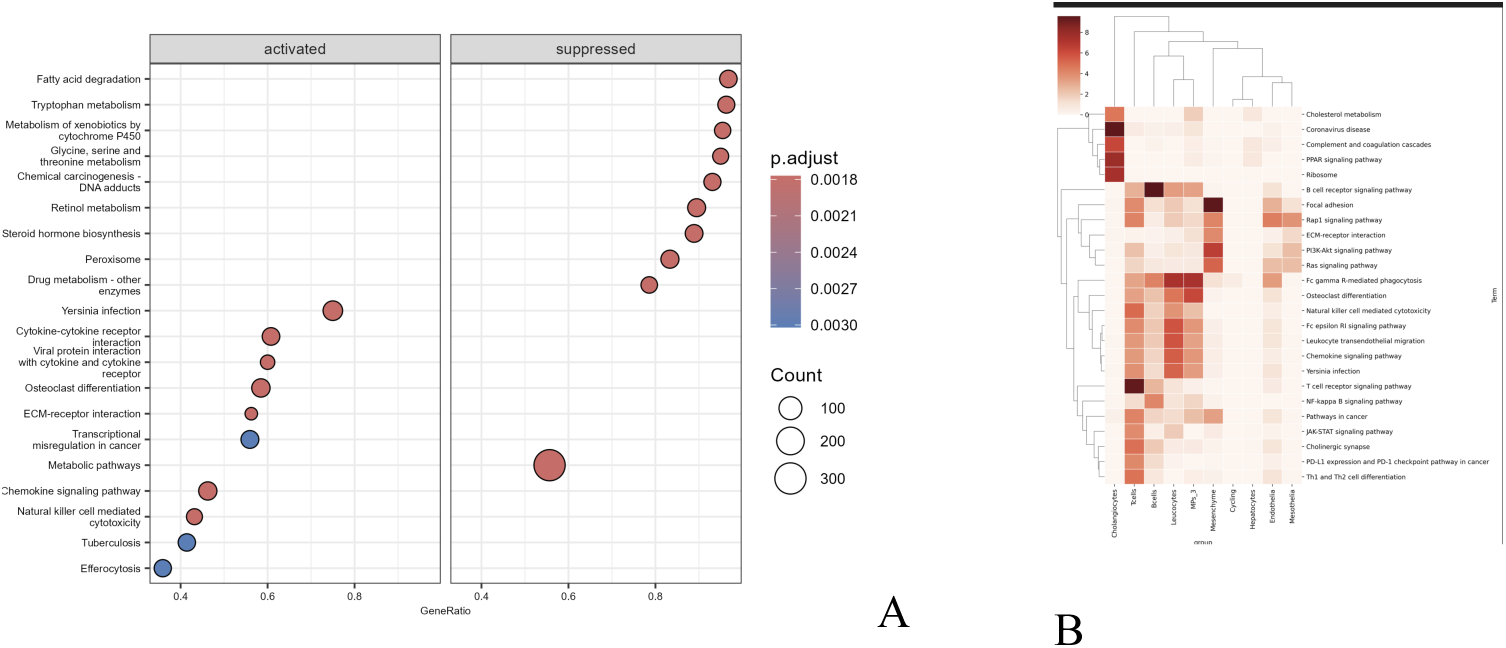
**A.** Differential Pathway Enrichment in MPs_3 (Volcano Plot). **B**. KEGG Pathway Heatmap Across Macrophage Subpopulations.

### 8. Macrophage Transition to a Reparative State Mediated by APOE–LRP1 Signaling

Cell-cell communication analysis revealed that apolipoprotein ligands (APOE, APOA1, APOB, APOC3) and binding proteins (HP, HPX) secreted by hepatocytes, cholangiocytes, endothelium, leukocytes, and mesenchyme converge on lipid-handling receptors (LRP1, LRP6, ABCA1, LDLR, SCARB1) expressed by the transitional macrophage subpopulation MPs_3. Parallel ECM cues, including FN1 and PLG activating ITGA9/ITGB1 integrins and APOE engaging SDC2, support matrix-lipid crosstalk, with FN1 emerging as a prominent ligand upregulated in peri-lesional zones by hepatic stellate and endothelial cells.

The MPs_3 population integrates these inputs through lipid scavenging, efferocytosis (via MARCO, MERTK, MSR1). This drives macrophage survival, cytoskeletal remodeling, and tissue repair, as evidenced by KEGG enrichments in chemokine signaling, Fcγ phagocytosis, and ECM-receptor interactions, contrasted with suppressed PPAR and fatty acid metabolism.

Pseudotime trajectory analysis depicted a progressive downregulation of CCR2-linked inflammatory features and upregulation of TREM2, SPP1, and cytoskeletal-ECM modules, resolving at repair-niche endpoints (branches 14–17). Longitudinal snRNA-seq mapping (0–300 h post-injury) pinpointed MPs_3 peak accumulation at 48 h, aligning with the activation-to-resolution transition.

Spatial integration of snRNA-seq with Visium transcriptomics (via cell2location, label transfer, and proximity-constrained L-R mapping) validated these dynamics in samples GSM6963518_pod24_a1 (24 h) and GSM6963526_pod48_d1 (48 h). At 48 h, APOE–LRP1 interactions enriched near peri-necrotic borders, yielding 19 APOE→LRP1.

Proximity mapping highlighted dense APOE–LRP1 and FN1–ITGA9 hotspots (orange-marked) at the necrotic border by 48 h, underscoring ITGA9/ITGB1’s role in FN1/tenascin-C/SPP1-mediated adhesion and FAK signaling for reparative retention. High-resolution overlays (e.g., segmenting from 2,599 to 1,009 cells) translated computational “nearby expression” into nanoscale protein-level evidence around MPs_3 niches, confirming MAPK1 involvement in downstream remodeling.

To validate the spatial distribution and temporal dynamics of reparative macrophages identified by single-cell transcriptomics, we performed histological analysis of cardiac tissue sections at defined timepoints post-injury. Early inflammatory infiltration was characterized by dense accumulation of mononuclear cells within damaged myocardium (panels A-B). Whole-heart cross-sections revealed preferential localization of macrophage-rich regions at the infarct border zone, forming a distinct transition between necrotic core and viable myocardium (panel C).

Higher-magnification views confirmed concentrated macrophage activity in critical repair zones (panel CD), consistent with the spatiotemporal enrichment patterns observed in our spatial transcriptomics analysis. Serial sections demonstrated progressive tissue remodeling and scar maturation (panels E-F), supporting the functional significance of the reparative macrophage population in orchestrating liver repair and maintaining ventricular structural integrity following ischemic injury (**Figure 11**).

**Figure 11.**
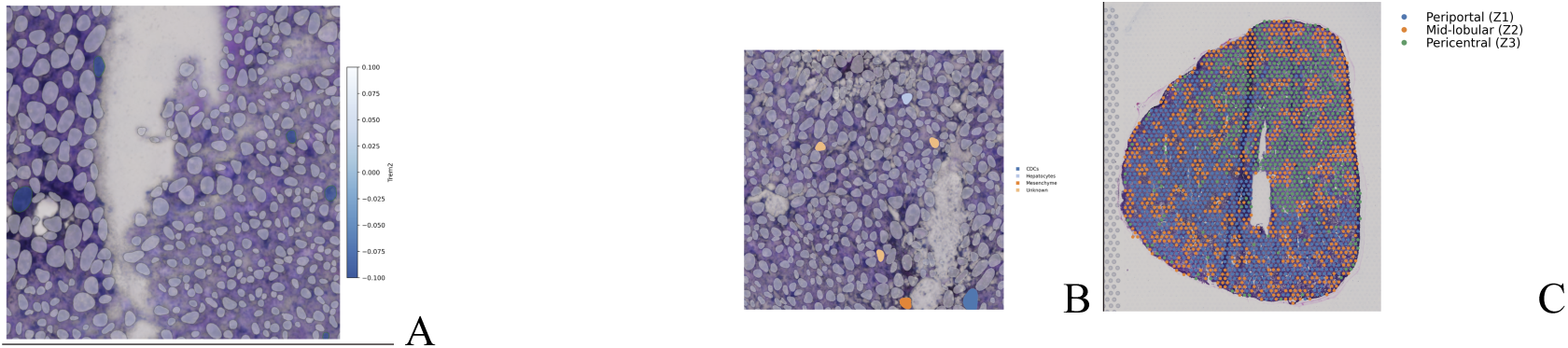
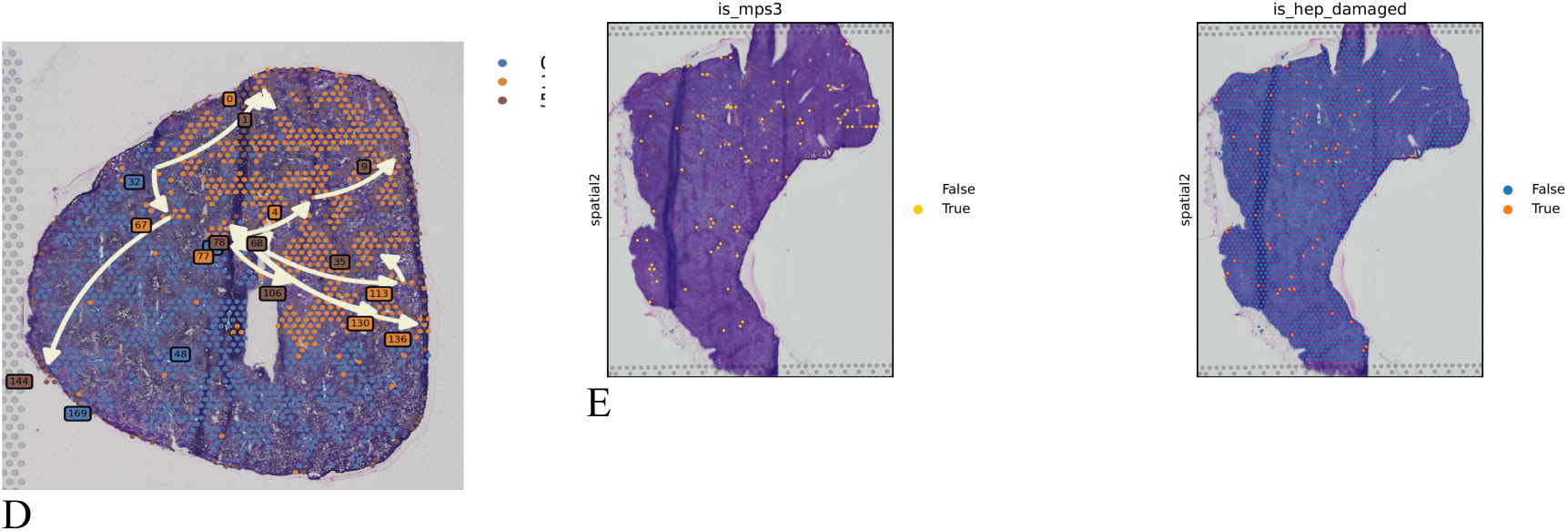
Figure: Histological validation of reparative macrophage localization in injured cardiac tissue. **(A)** High-magnification view of inflammatory cell infiltrate in damaged myocardium at early timepoint (H&E staining), showing dense cellular accumulation. **(B)** Representative section demonstrating macrophage distribution within the infarct border zone. **(C)** Low-magnification cardiac cross-section with outlined infarct region (dashed line) showing spatial organization of immune cell infiltration relative to viable myocardium. **(D)** Composite cardiac section with highlighted regions of interest (yellow dashed boxes) marking areas of concentrated macrophage activity at the infarct border and within the necrotic core. **(E)** Serial sections of whole heart showing the temporal and spatial evolution of macrophage recruitment and tissue remodeling, with blue coloration indicating fibrotic scar formation and preservation of ventricular architecture.

Spatial profiling confirms APOE upregulation in parenchymal zones interfacing necrotic lesions (mean expression +0.3), decreasing toward central veins (−0.2), supporting its role as a hepatocyte-derived cue for macrophage efferocytosis and repair initiation at 48 h. Figure 18.B. Spatial Co-Localization of Hepatocytes and MPs_3 Macrophages in Peri-Necrotic Zones.

Overlay of cell-type deconvolution in a 48 h Visium section, with hepatocytes (blue) predominant in parenchymal regions and MPs_3 macrophages (orange) enriched at injury interfaces (white: necrotic voids). This zonal patterning underscores APOE-LRP1 signaling hotspots, facilitating lipid transfer and efferocytosis during resolution. C. Zonal Architecture of Hepatic Lobules in Spatial Transcriptomics.

Color-coded zonation map of a liver section, delineating periportal (blue), midlobular (orange), and pericentral (green) regions based on transcriptomic gradients. MPs_3 macrophages (darker orange hotspots) preferentially localize to midlobular-pericentral interfaces near necrotic zones, aligning with elevated APOE–LRP1 and FN1–ITGA9 signaling for targeted repair. Figure 18 D. Binary Spatial Maps of MPs_3 Macrophage and Damaged Hepatocyte Localization Left: Probability map of MPs_3 enrichment (magenta: high-confidence true spots; purple: false/low; yellow/orange: trust scores), revealing discrete peri-lesional patches. Right: Binary map of hepatocyte damage (red: true damaged spots; blue: false undamaged; orange: validated true). Co-enrichment at injury borders (48 h section) highlights MPs_3 targeting of necrotic zones for efferocytosis and repair. Figure 18 E. Temporal Cascade of Macrophage Activation and Repair Signaling

## Discussion

We present TRIO-AI, a hybrid computational framework that synergistically integrates Temporal Graph Neural Networks, Neural Ordinary Differential Equations, and Time-Variational Autoencoders to provide unprecedented resolution in single-cell trajectory inference. This multi-level approach addresses fundamental limitations of existing methods by combining discrete branching topology detection, continuous state flow modeling, and density-aware transitional state identification.

Application of TRIO-AI to liver ischemia-reperfusion injury models revealed a novel transitional macrophage population (MPs_3) that bridges inflammatory and reparative phases. This population exhibits a coordinated transcriptional program integrating lipid scavenging, efferocytosis, and ECM engagement, orchestrated through convergent APOE-LRP1 and FN1-ITGA9 signaling axes. The identification of MPs_3 provides new mechanistic insights into macrophage-mediated tissue repair and offers potential therapeutic targets for modulating injury resolution.

### Technical Advances Over Existing Methods

TRIO-AI demonstrates multiple technical advantages compared to conventional trajectory inference approaches. First, the Temporal GNN layer effectively incorporates RNA velocity directionality and experimental timepoint information into graph construction, enabling discovery of biologically meaningful branching patterns that align with actual temporal progression. This represents a substantial improvement over pseudotime-based methods that struggle to capture true temporal dynamics.

Second, the Neural ODE layer provides continuous modeling of state transitions, complementing discrete graph-based branching detection. This dual representation captures both major lineage bifurcations and subtle off-path transitions, with particular sensitivity to rare or alternative fates missed by cluster-level methods. The ability to operate at both latent-time bin and cluster granularities provides comprehensive trajectory mapping from fine-grained intermediate states to major phenotypic transitions.

Third, the Time-VAE density layer quantifies sampling probability across the temporal manifold, systematically identifying under-sampled transitional regions that represent unstable or rapidly transitioning cell states. This probabilistic framework provides rigorous criteria for distinguishing stable attractor states from transient intermediates, addressing a critical gap in traditional clustering approaches that treat all populations equivalently.

### Biological Insights into Macrophage Dynamics

The discovery and characterization of MPs_3 provides several important biological insights. First, the temporal dynamics of this population—peaking at 48 hours post-injury followed by rapid decline—suggest it represents a critical checkpoint in the transition from inflammatory to reparative macrophage programming. The transient nature of MPs_3 implies active regulation rather than a stable differentiation endpoint.

Second, the coordinated lipid-handling signature of MPs_3, featuring multiple apolipoprotein receptors (LRP1, LRP6, ABCA1, LDLR, SCARB1), indicates metabolic reprogramming as a key feature of the inflammatory-to-reparative transition. This is consistent with emerging evidence that lipid metabolism regulates macrophage polarization and that dysregulated lipid handling contributes to pathological inflammation.

Third, the spatial localization of MPs_3 to peri-necrotic borders with enriched ligand-receptor co-localization at 48 hours demonstrates anatomical organization of the reparative response. The discrete, speckled distribution pattern suggests targeted recruitment mechanisms responding to local tissue damage signals rather than systemic activation.

### Therapeutic Implications

The identification of MPs_3 and its defining molecular programs suggests potential therapeutic strategies for modulating liver injury resolution. The APOE-LRP1 axis represents an attractive target, as enhancing this pathway could accelerate the inflammatory-to-reparative transition. Similarly, the FN1-ITGA9 interaction provides a therapeutic handle for manipulating macrophage localization and retention in injury sites.

More broadly, the surface-accessible nature of key MPs_3 markers (LRP1, ITGA9) provides practical handles for validation studies using proximity ligation assays, immunofluorescence, flow cytometry, and functional perturbation experiments. Blocking studies with LRP1 or ITGA9 antibodies, or rescue experiments with recombinant APOE or FN1, could establish causal roles for these interactions in macrophage state transitions.

### Limitations and Future Directions

While TRIO-AI demonstrates substantial advances, several limitations warrant consideration. First, the framework requires actual experimental timepoints for optimal performance, limiting application to purely cross-sectional datasets. However, integration with RNA velocity can partially compensate for absent temporal information.

Second, computational complexity increases substantially compared to conventional methods, particularly for large datasets. Future optimizations including GPU acceleration and algorithmic refinements could improve scalability. Third, interpretation of Neural ODE trajectories requires biological context and domain expertise, as not all computationally identified paths necessarily represent biologically realized transitions.

Fourth, while spatial validation strengthens MPs_3 identification, experimental functional validation remains essential. Future work should include lineage tracing, perturbation experiments, and clinical correlation studies to establish the causal role of MPs_3 in injury resolution and assess translatability to human disease.

Finally, extension of TRIO-AI to additional biological systems and experimental contexts will establish generalizability. Applications to developmental, regenerative, and disease systems beyond liver injury could reveal whether the framework successfully captures transitional dynamics across diverse biological processes.

### Conclusions

TRIO-AI represents a significant methodological advance in single-cell trajectory inference, providing integrated discrete-continuous modeling with explicit temporal awareness and density-based transitional state detection. The framework successfully identified a novel macrophage population bridging inflammatory and reparative phases in liver injury, characterized by coordinated lipid-handling, efferocytosis, and ECM engagement programs orchestrated through APOE-LRP1 and FN1-ITGA9 signaling axes.

This work demonstrates the power of hybrid computational approaches that synergistically combine complementary algorithmic strategies. By integrating graph neural networks, differential equations, and variational autoencoders, TRIO-AI achieves sensitivity to rare and transitional populations while maintaining robust reconstruction of major lineage trajectories. This balance between comprehensive coverage and fine-grained resolution addresses a longstanding challenge in trajectory inference.

Beyond specific biological insights into liver injury and macrophage dynamics, TRIO-AI provides a generalizable platform for dissecting cellular state transitions in complex biological systems. The framework’s ability to detect transient, rare, and off-path populations positions it as a valuable tool for discovering novel cell states and regulatory mechanisms across diverse experimental contexts.

## Notes

### Competing Interest Statement

The authors have declared no competing interest.

### Funding

- Cancer Prevention Research Institute of Texas (CPRIT)

## Reference

1. Cannistrà M, Ruggiero M, Zullo A, Gallelli G, Serafini S, Maria M, Naso A, Grande R, Serra R, Nardo B. Hepatic ischemia reperfusion injury: A systematic review of literature and the role of current drugs and biomarkers. Int J Surg. 2016 Sep;33 Suppl 1:S57–70. doi: 10.1016/j.ijsu.2016.05.050. Epub 2016 May 30. PMID: 27255130.

2. Zhang M, Liu Q, Meng H, Duan H, Liu X, Wu J, Gao F, Wang S, Tan R, Yuan J. Ischemia-reperfusion injury: molecular mechanisms and therapeutic targets. Signal Transduct Target Ther. 2024 Jan 8;9(1):12. doi: 10.1038/s41392-023-01688-x. PMID: 38185705; PMCID: PMC10772178.

3. Ma X, Qiu J, Zou S, Tan L, Miao T. The role of macrophages in liver fibrosis: composition, heterogeneity, and therapeutic strategies. Front Immunol. 2024 Nov 20;15:1494250. doi: 10.3389/fimmu.2024.1494250. PMID: 39635524; PMCID: PMC11616179.

4. Wang W, Li S, Liu Y, Ding X, Yang Y, Chen S, Cao J, Tacke F, Dong W, Lan T. Macrophage heterogeneity in liver fibrosis. Front Immunol. 2025 Sep 4;16:1639455. doi: 10.3389/fimmu.2025.1639455. PMID: 40977736; PMCID: PMC12443557.

5. Yang Y, Wang M, Qiu X, Yang R, Yao C. Single-cell RNA sequencing: new insights for pulmonary endothelial cells. Front Cell Dev Biol. 2025 Jun 17;13:1576067. doi: 10.3389/fcell.2025.1576067. PMID: 40599812; PMCID: PMC12209310.

6. Yoshihara K, Ito K, Kimura T, Yamamoto Y, Urabe F. Single-cell RNA sequencing and spatial transcriptome analysis in bladder cancer: Current status and future perspectives. Bladder Cancer. 2025 Feb 21;11(1):23523735251322017. doi: 10.1177/23523735251322017. PMID: 40034247; PMCID: PMC11864234.

7. Hora S, Wuestefeld T. Liver Injury and Regeneration: Current Understanding, New Approaches, and Future Perspectives. Cells. 2023 Aug 22;12(17):2129. doi: 10.3390/cells12172129. PMID: 37681858; PMCID: PMC10486351.

8. Huang R, Zhang X, Gracia-Sancho J, Xie WF. Liver regeneration: Cellular origin and molecular mechanisms. Liver Int. 2022 Jul;42(7):1486–1495. doi: 10.1111/liv.15174. Epub 2022 Feb 24. PMID: 35107210.

9. Xu X, Su J, Zhu R, Li K, Zhao X, Fan J, Mao F. From morphology to single-cell molecules: high-resolution 3D histology in biomedicine. Mol Cancer. 2025 Mar 3;24(1):63. doi: 10.1186/s12943-025-02240-x. PMID: 40033282; PMCID: PMC11874780.

10. Kitaoka, Y.; Uchihashi, T.; Kawata, S.; Nishiura, A.; Yamamoto, T.; Hiraoka, S.-i.; Yokota, Y.; Isomura, E.T.; Kogo, M.; Tanaka, S.;, et al. Role and Potential of Artificial Intelligence in Biomarker Discovery and Development of Treatment Strategies for Amyotrophic Lateral Sclerosis. Int. J. Mol. Sci. 2025, 26, 4346. 10.3390/ijms2609434

11. Zhong J, Gao RR, Zhang X, Yang JX, Liu Y, Ma J, Chen Q. Dissecting endothelial cell heterogeneity with new tools. Cell Regen. 2025 Mar 23;14(1):10. doi: 10.1186/s13619-025-00223-3. PMID: 40121354; PMCID: PMC11929667.

12. Cole Trapnell*, Davide Cacchiarelli*, Jonna Grimsby, Prapti Pokharel, Shuqiang Li, Michael Morse, Niall J. Lennon, Kenneth J. Livak, Tarjei S. Mikkelsen, and John L. Rinn.The dynamics and regulators of cell fate decisions are revealed by pseudotemporal ordering of single cells. Nature Biotechnology, 2014

13. Street K, Risso D, Fletcher RB, Das D, Ngai J, Yosef N, Purdom E, Dudoit S. Slingshot: cell lineage and pseudotime inference for single-cell transcriptomics. BMC Genomics. 2018 Jun 19;19(1):477. doi: 10.1186/s12864-018-4772-0. PMID: 29914354; PMCID: PMC6007078.

14. Volker Bergen, Marius Lange, Stefan Peidli, F. Alexander Wolf and Fabian J. Theis Generalizing RNA velocity to transient cell states through dynamical modeling Nature Biotechnology 38, 1408–1414 (2020)

15. Lange M, Bergen V, Klein M, Setty M, Reuter B, Bakhti M, Lickert H, Ansari M, Schniering J, Schiller HB, Pe’er D, Theis FJ. CellRank for directed single-cell fate mapping. Nat Methods. 2022 Feb;19(2):159–170. doi: 10.1038/s41592-021-01346-6. Epub 2022 Jan 13. PMID: 35027767; PMCID: PMC8828480.

16. Tran TN, Bader GD. Tempora: Cell trajectory inference using time-series single-cell RNA sequencing data. PLoS Comput Biol. 2020 Sep 9;16(9):e1008205. doi: 10.1371/journal.pcbi.1008205. PMID: 32903255; PMCID: PMC7505465.

17. Tong A, Huang J, Wolf G, van Dijk D, Krishnaswamy S. TrajectoryNet: A Dynamic Optimal Transport Network for Modeling Cellular Dynamics. Proc Mach Learn Res. 2020 Jul;119:9526–9536. PMID: 34337419; PMCID: PMC8320749.

18. Traag VA, Waltman L, van Eck NJ. From Louvain to Leiden: guaranteeing well-connected communities. Sci Rep. 2019 Mar 26;9(1):5233. doi: 10.1038/s41598-019-41695-z. PMID: 30914743; PMCID: PMC6435756.

19. Seth S, Mallik S, Bhadra T, Zhao Z. Dimensionality Reduction and Louvain Agglomerative Hierarchical Clustering for Cluster-Specified Frequent Biomarker Discovery in Single-Cell Sequencing Data. Front Genet. 2022 Feb 7;13:828479. doi: 10.3389/fgene.2022.828479. PMID: 35198011; PMCID: PMC8859265.

20. Matchett KP, Wilson-Kanamori JR, Portman JR, Kapourani CA et al. Multimodal decoding of human liver regeneration. Nature 2024 Jun;630(8015):158–165. PMID: 38693268

21. Wen Z, Fang Y, Wei P, Liu F, Chen Z, Wu M. Temporal and Heterogeneous Graph Neural Network for Remaining Useful Life Prediction. IEEE Trans Neural Netw Learn Syst. 2025 Nov;36(11):19748–19761. doi: 10.1109/TNNLS.2025.3592788. PMID: 40748812.

22. u, W., Gao, F., Lou, X., et al. Discovering latent node Information by graph attention network. Sci Rep 11, 6967 (2021). 10.1038/s41598-021-85826-x

23. Jan Z, Shabir M, Farman H, Rahman A, M Nasralla M. Deep learning based semantic segmentation of leukemia effected white blood cell. PLoS One. 2025 May 8;20(5):e0320596. doi: 10.1371/journal.pone.0320596. PMID: 40338981; PMCID: PMC12061112.

